# Facilitating genome annotation using ANNEXA and long-read RNA sequencing

**DOI:** 10.1101/2025.04.16.648718

**Authors:** Nicolaï Hoffmann, Aurore Besson, Edouard Cadieu, Matthias Lorthiois, Victor Le Bars, Armel Houel, Christophe Hitte, Catherine André, Benoit Hédan, Thomas Derrien

## Abstract

With the advent of complete genome assemblies, genome annotation has become essential for the functional interpretation of genomic data. Long-read RNA sequencing (LR-RNAseq) technologies have significantly improved transcriptome annotation by enabling full-length transcript reconstruction for both coding and non-coding RNAs. However, challenges such as transcript fragmentation and incomplete isoform representation persist, highlighting the need for robust quality control (QC) strategies. This study presents ANNEXA, a pipeline designed to enhance genome annotation using LR-RNAseq data while also providing QC for reconstructed genes and transcripts. ANNEXA integrates two transcriptome reconstruction tools, StringTie2 and Bambu, applying stringent filtering criteria to improve annotation accuracy. It also incorporates deep learning models to evaluate transcription start sites (TSSs) and employs the tool FEELnc for the systematic annotation of long non-coding RNAs (lncRNAs). Additionally, the pipeline offers intuitive visualisations for comparative analyses of coding and non-coding repertoires. Benchmarking against multiple reference annotations revealed distinct patterns of sensitivity and precision for both known and novel genes and transcripts and mRNAs and lncRNAs. To demonstrate its utility, ANNEXA was applied in a cross-species study involving LR-RNAseq of two human and eight canine cancer cell lines of mucosal melanoma. The pipeline successfully identified novel genes and transcripts across species, expanding the catalog of protein-coding and lncRNA annotations in both species. Implemented in Nextflow for scalability and reproducibility, ANNEXA is available as an open-source tool: https://github.com/IGDRion/ANNEXA.

## Introduction

With the increasing availability of entire genome assemblies, i.e. telomere-to-telomere (T2T) (Nurk et al., 2022), one challenge in genome research is to move from improving genome completeness to refining genome annotation. High-quality genome sequences now enable a more precise characterisation of genes and transcripts, especially in repetitive regions of the genomes, making transcriptome-based annotation a critical step in the functional interpretability of the genomes (Nurk et al., 2022). To this end, RNA sequencing (RNAseq) plays a central role in this process by providing direct transcript-level evidence, essential for defining gene structures, alternative splicing events, and non-coding RNA repertoires.

While short-read RNAseq (SR-RNASeq) has shown some limitations to reconstruct full-length transcripts (Steijger et al., 2013), long-read RNA sequencing (LR-RNAseq), provided by platforms such as Pacific Biosciences (PacBio) and Oxford Nanopore Technologies (ONT), has significantly advanced transcriptome annotation by providing reads that span repeats and also by allowing direct connectivity between distant exons of the same isoform (Conesa et al., 2016). However, despite these advantages, LR-RNAseq-based transcriptome reconstruction still remains prone to artifacts such as truncated transcript models and partial isoform representation (Sessegolo et al., 2019). To ensure accurate genome annotation, robust quality control (QC) and filtering strategies are needed to evaluate and refine transcriptome reconstructions.

Several tools have been developed to assemble long reads from LR-RNAseq data. A recent benchmark study from the LRGASP consortium has compared fourteen transcriptome reconstruction and quantification tools (Pardo-Palacios et al., 2024b) and showed that choosing the best program depends on the biological context of the study and the completeness of the reference annotation. Among the benchmarked tools, Bambu (Chen et al., 2023) and StringTie (Kovaka et al., 2019) consistently demonstrated strong performance across multiple metrics. However, one limitation of this benchmark is that it did not explicitly assess tool performance with respect to RNA biotypes, particularly in distinguishing between protein-coding transcripts (mRNAs) and long non-coding RNAs (lncRNAs). Given the complexity and heterogeneity of transcriptomes, this distinction could be important since different RNA biotypes may vary in expression levels, structural features, and evolutionary conservation, all of which can impact reconstruction accuracy. This consideration is especially important in light of the substantial expansion of the human lncRNA catalogue in recent Gencode releases (Kaur et al., 2024), underscoring the need for tools that can accurately reconstruct and annotate both known and novel transcripts across diverse RNA classes.

A number of pipelines dedicated to long-read RNAseq analysis and annotation extension integrate some of the tools benchmarked in LRGASP but present certain limitations. For example, nfcore nanoseq (Chen et al., 2025) supports transcriptome reconstruction using Bambu or StringTie but it is restricted to Oxford Nanopore data and does not annotate long non-coding RNAs. TAGADA (Kurylo et al., 2023) and annotate_my_genomes (Farkas et al., 2022) offer lncRNA detection but rely exclusively on StringTie for transcript assembly and short-read RNASeq for TAGADA. To overcome these limitations, we developed ANNEXA, a Nextflow-based pipeline specifically designed to extend reference annotations from long-read RNAseq data while also assessing the quality of newly identified transcript models. ANNEXA integrates two transcriptome reconstruction tools, Bambu (Chen et al., 2023) and StringTie (Kovaka et al., 2019), and provides users with stringent filters to improve annotation accuracy. It incorporates a deep learning strategy to potentially remove truncated transcript models by evaluating transcription start sites (TSSs) of all novel transcripts. ANNEXA is also designed to systematically annotate and evaluate the annotation of lncRNAs by incorporating the FEELnc program (Wucher et al., 2017). It provides intuitive visual representations of the annotation, facilitating comparative analysis of coding and non-coding repertoires. To illustrate the usability of ANNEXA in a comparative oncology project, we sequenced two human and eight canine cancer cell lines from mucosal melanomas, histiocytic sarcomas and osteosarcoma using ONT direct cDNA sequencing, and identified novel human and canine genes/transcripts, some of which being conserved in the two species.

By implementing a structured framework for quality control and annotation, ANNEXA enhances the reliability of long-read transcriptomics, ensuring more comprehensive and biologically meaningful extended genome annotations as a prerequisite for further functional studies.

## Methods

### Overview of ANNEXA

ANNEXA is a pipeline that extends user-provided reference annotations with novel genes and transcript isoforms from long-read sequencing data. It only uses three parameter files: a reference genome, a reference annotation and a text file listing the paths to BAM file(s). (**Fig. 1.A**). Unlike technology-specific pipelines such as nf-core/isoseq (Guizard et al., 2023), which is tailored for PacBio data, ANNEXA is compatible with both Nanopore and PacBio RNA sequencing technologies. The pipeline is organised into four main modules, described below:

1. Transcriptome reconstruction and quantification
2. Coding potential evaluation and transcript classification
3. Transcript filtering and full-length assessment
4. Quality Control

**Figure 1.**
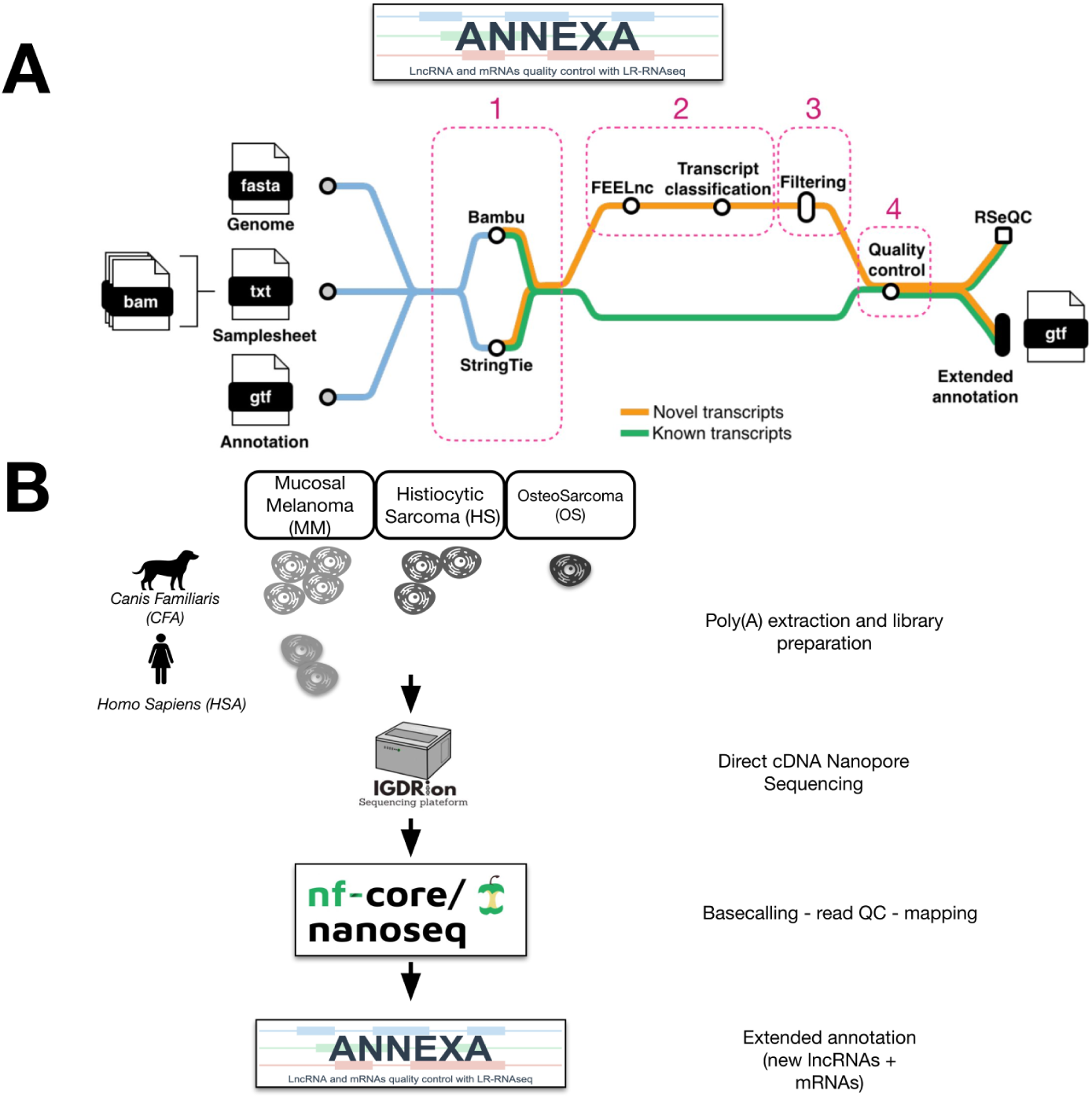
**A**- Metromap of ANNEXA. The four main modules are circlized (in pink) with module 1=Transcriptome reconstruction and quantification, 2=Coding potential evaluation and transcript classification, 3=Transcript filtering and full-length assessment and 4=Quality Control **B**- Experimental design with species-specific input cancer cell lines and experimental and computational analyses.

### Transcriptome reconstruction and quantification

As recently demonstrated by the LRGASP consortium, the choice of the bioinformatic tool to model known and novel genes/transcripts depends on the biological context, and more particularly, on the expected levels of annotation precision (controlling the number of novel false-positive transcripts) and sensitivity/recall (controlling the number of novel false-negative transcripts). Ideally, users should be able to select the appropriate tool based on the expected number of novel transcripts and the quality or completeness of the reference genome and annotation. In this study, transcriptome completeness is defined as the total number of genes and transcripts catalogued in a given annotation. To support this, ANNEXA enables the reconstruction and quantification of both known and novel transcripts using two distinct tools: Bambu (Chen et al., 2023) and StringTie2 (Kovaka et al., 2019). When multiple input files are provided, ANNEXA pools together all samples to produce a single extended annotation. With StringTie, a first step is performed where each sample is processed separately, producing a GTF file specific to that sample. In a second step, all GTF files are merged together with the reference annotation using the `-merge` option from StringTie, producing a unified, non-redundant transcript set. With Bambu, it takes all samples together to process reads, and uses a machine learning approach to define a TPS (Transcript Probability Score) corresponding to the probability that a transcript is valid in at least one sample.

In ANNEXA, several options are available for Bambu, including the ability to adjust the Novel Discovery Rate (NDR) threshold, a key metric derived from the TPS that balances sensitivity and precision (Chen et al., 2023) across multiple samples. The NDR estimates an upper limit on the false discovery rate under the assumption of complete reference annotations (e.g., NDR=0.1 indicates that at least 90% of transcripts with similar or higher scores are annotated (Chen et al., 2023)). ANNEXA’s users can enhance Bambu sensitivity by increasing the default recommended NDR threshold with the `–-bambu_threshold` option, and include single-exon transcripts with the `–-bambu_singleexon` option. Alternatively, users can choose to use the NDR threshold recommended by Bambu by enabling the `–-bambu_rec_ndr` option, which replicates Bambu’s default behaviour.

For StringTie2, ANNEXA uses the long reads assembly mode (-L) for the reconstruction step on each individual sample and the -merge option to produce a unified transcript set.

Gene and transcript quantifications are performed internally by both tools. Bambu directly produces two raw count tables for gene and transcripts, while for Stringtie the normalised counts are converted to raw counts using the `extractGeneExpression` function from the IsoformSwitch-AnalyzeR program (Vitting-Seerup and Sandelin, 2019) and all the expression tables are made available by default in the final output of ANNEXA.

At the end of this module, ANNEXA integrates all genes and transcripts from the reference annotation along with the novel transcript models from StringTie2 or Bambu into an unfiltered extended annotation.

### Coding potential and transcript classification

Among the newly assembled transcripts, it is essential to annotate different RNA categories, particularly distinguishing protein-coding transcripts (mRNAs) from long non-coding RNAs (lncRNAs). To achieve this, ANNEXA integrates the FEELnc program (Wucher et al., 2017) to predict the coding potential of all novel transcripts from novel genes (yellow line, **Fig. 1.A**). To accelerate the FEELnc process, we previously demonstrated that training it on a subset of known mRNAs and lncRNAs from the reference annotation preserves high predictive performance (Wucher et al., 2017), thereby allowing ANNEXA to use FEELnc with a, by default, random subset of 3,000 reference lncRNAs and mRNAs. However, in the instance where the user’s reference annotation lacks lncRNAs or contains less than 3,000 transcripts, we have implemented the `–-feelnc_mrna` and `–-feelnc_lncrna` options which allow them to define a number of transcripts to use.

To assess the potential impact of genomic alterations (mutations) on newly identified protein-coding transcripts, ANNEXA employs TransDecoder (https://github.com/TransDecoder/TransDecoder) using the `TransDecoder.LongOrfs` module to extract longest ORFs and `TransDecoder.Predict` with the `–single_best_only` option to predict coding regions, ensuring that only the single best ORF per transcript is retained while reporting the corresponding CDS (Coding Determining Sequence) information in the final GTF file. FEELnc is applied to all novel transcripts, both from novel and known genes, while TransDecoder is applied to all novel transcripts classified as protein-coding by FEELnc. Additionally, to classify novel transcripts with respect to the input reference annotation, ANNEXA uses GffCompare (Pertea and Pertea, 2020), incorporating this information under the `class_code` attribute in the extended GTF. This enables users to efficiently extract specific transcript classes, such as `class` `k` corresponding to alternative isoforms extending known genes at the 5’ or 3’ ends or `class` `x`, corresponding to exonic antisense transcripts (often classified as antisense lncRNA biotype).

### Transcript filtering and full-length assessment

While long-read RNA sequencing (LR-RNAseq) has significantly improved transcriptome reconstruction by capturing full-length isoforms, it still exhibits biases that lead to fragmented transcript models, particularly at the 5’ end *i.e.* Transcription Start Sites (TSSs) (Tardaguila et al., 2018). Here, transcript full-lengthness refers to the extent to which transcript models include the complete sequence until the transcription start site (TSS).

To refine the extended annotation file and exclude novel incomplete transcripts, ANNEXA employs two complementary filtering strategies: (i) the Novel Discovery Rate (NDR) cutoff, derived from the Bambu tool, which evaluates the confidence of novel transcript predictions and (ii) the TransforKmer cutoff which assesses the likelihood of novel transcript transcription start sites (TSSs) using a species-specific deep learning model. The TransforKmer model (TFK) is pre-trained with DNABERT (Ji et al., 2021), a transformer-based architecture designed for genomic sequence analysis, and further fine-tuned on a classification task using labeled TSSs from the reference annotations (Karollus et al., 2024). By default, ANNEXA applies a conservative NDR cutoff to retain high-confidence novel transcripts while filtering out likely incomplete models. The TFK filtering step is subsequently applied to further assess the likelihood that novel transcript models contain a valid transcription start site (TSS). Both the TFK-filtered and unfiltered annotations are retained, allowing users to balance transcript discovery and transcript full-lengthness according to their specific needs. As demonstrated in our stability analysis (**Supp. Fig. 1**), users can also adjust the TFK threshold to increase sensitivity, at the cost of retaining more novel transcripts that may be less likely to be full-length.

TransforKmers performance was evaluated using annotated transcript TSSs as positive examples and two types of negative controls: shuffled sequences preserving nucleotide composition and fully random sequences. Model outputs show clear separation between true TSSs and negative controls (**Supp. Fig. 2.A**). In addition, TFK classification accuracy was stratified by transcript biotype and transcript support level (TSL) in human, revealing a correlation between TFK accuracy and well-supported transcripts (**Supp. Fig. 2.B**).

**Figure 2.**
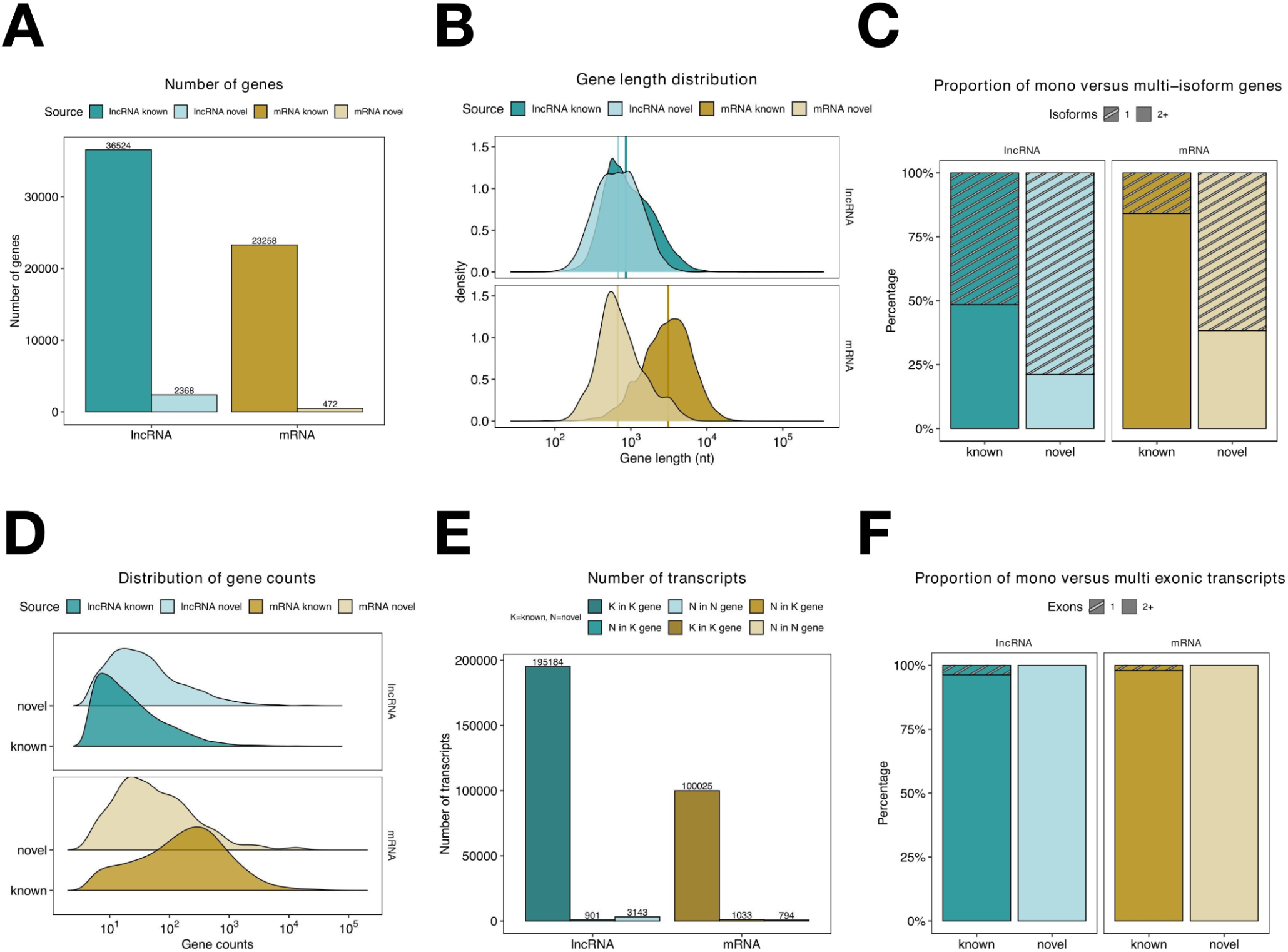
Example of an automatic QC report from ANNEXA. This report was generated from the analysis of two human mucosal melanoma samples using Bambu with the Gencode annotation (the full PDF can be found in the GitHub repository of ANNEXA and a detailed explanations of each panel in the dedicated wiki page: https://github.com/IGDRion/ANNEXA/wiki/ANNEXA-wiki). **A**- Number of known and novel genes. **B**- Gene length distribution. **C**- Proportion of gene mono-versus multi-isoform(s). **D**- Distribution of gene read counts. **E**- Number of transcripts in known (K) and novel (N) genes. **F**- Proportion of transcript mono/single- versus multi-exonic.

To adjust the stringency of the filtering process, users can choose between two modes: (i) the union of the two filters (filtering `operation` `=` `union`), which retains transcripts passing at least one filter, or (ii) the intersection (filtering `operation` `=` `intersection`), which retains only transcripts passing both filters (**Supp. Fig. 3**). At the end of this module, ANNEXA outputs an additional more stringent filtered annotation. Because ANNEXA provides both a full and filtered annotations as output, researchers can flexibly tailor their analysis, prioritising either sensitivity or precision, based on their specific research goals and biological questions.

**Figure 3.**
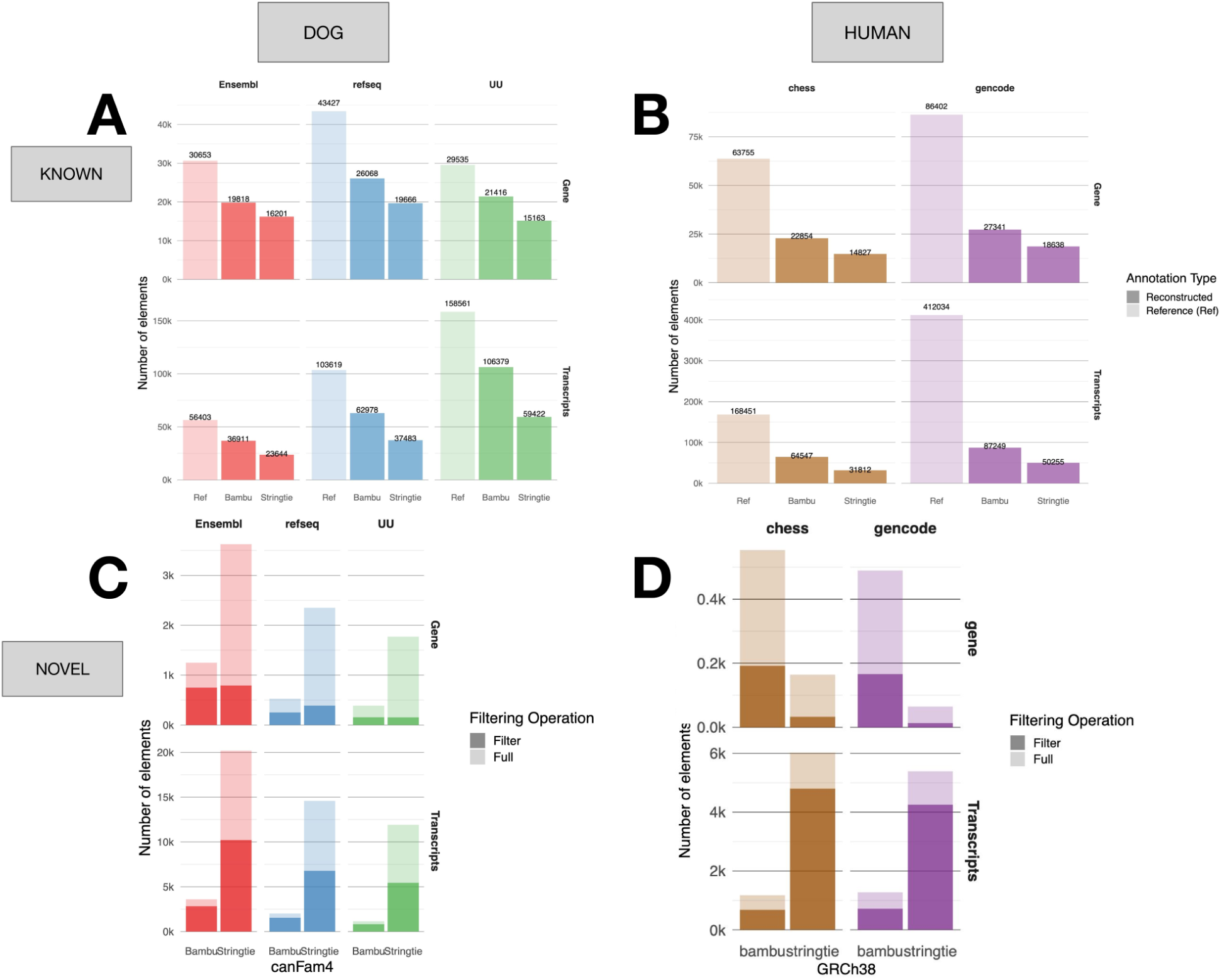
Benchmark analysis between Bambu and StringTie for dog (left panels) and human (right panels) for known (top panel) and novel (bottom panel) genes (upper part of each panel) and transcripts (bottom part). For known elements (top), the number of genes/transcripts from the reference annotation (transparent bars) is separated from reconstructed genes/transcripts by Bambu and StringTie (solid bars). **A**- Number of known canine genes/transcripts retrieved from three reference annotations (Ensembl - red, ref-seq - blue and UnivUppsala - green). **B**- Number of known human genes/transcripts retrieved from two reference annotations (Chess - brown and Gencode - purple). **C**- Number of novel dog genes/transcripts. Full set of elements (light bar) are separated from ANNEXA’s filtered (intersection) set (dark bar). **D**- Number of novel human genes/transcripts detected on canonical chromosomes (chr1-22, X, Y and M). Full set of elements (light bar) are separated from ANNEXA’s filtered (intersection) set (dark bar).

### Quality Control

The two resulting annotation files (*full* and *filtered*) may contain thousands of novel isoforms, including both mRNAs and lncRNAs, which require thorough inspection for final quality control (QC). Inspired by the SQANTI tool (Pardo-Palacios et al., 2024a), ANNEXA computes multiple features by comparing known (*i.e.*, matching the reference annotation) and novel (*i.e.*, reconstructed by Bambu or StringTie) genes, transcripts, and exons, collectively referred to as annotated elements (AE). The QC metrics can be broadly categorised into two groups:

- Structural metrics of AEs: including the number of known and novel AEs, AE length distribution, proportion of single- versus multi-isoform genes (at the gene level), and proportion of single- versus multi-exonic transcripts (at the transcript level).
- Quantification-related metrics: such as the distribution of gene read counts across input samples or the breadth of expression of the AEs.

Additionally, an optional QC feature assesses gene body coverage by sample reads using the RSeQC pipeline (Wang et al., 2012), allowing the identification of potential 5’ or 3’ read-coverage biases across known and novel genes.

All QC indicators are generated as CSV files, which serve as input for ANNEXA to compile and visualise the data in a comprehensive final QC report. This report contains information at three AE levels (gene, transcript and exon) and summarises key metrics, such as the number of novel elements, their length distribution, and expression counts as illustrated in **Fig. 2**.

### ANNEXA implementation

ANNEXA is implemented in Nextflow (Di Tommaso et al., 2017) and integrates scripts written in multiple languages, including R, Python and Bash. The Nextflow framework is designed to facilitate the development of reproducible, scalable, and portable analysis workflows. ANNEXA can use both Docker and Singularity containers and has been successfully executed on standalone computers and high-performance computing clusters using SGE or SLURM. The project is open-source, with its code accessible on GitHub: https://github.com/IGDRion/ANNEXA/.

### Long-Read Sequencing protocol of cancer cell lines

#### Experimental methods

Ten LR-RNAseq experiments were conducted (**Fig. 1.B**) with eight canine cancer cell lines from three canine cancer types: Mucosal Melanoma (MM, n=4), Histiocytic Sarcoma (HS, n=3) and Osteosarcoma (OS, n=1) and two human cancer cell lines also originating from MM patients. For all samples, RNA was extracted from *∼*15 million cells using the NucleoSpin RNA kit (Macherey-Nagel). Library preparation was performed according to manufacturer’s protocol (Oxford Nanopore Technologies, ONT) with the direct cDNA Sequencing Kit (SQK-DCS109). Sequencing was done using MinION Flow Cells (FLO-MIN106D) with GridION device from the IGDRION platform (https://igdr.univ-rennes.fr/igdrion).

#### Computational methods

Basecalling of `fast5` files was done with guppy (version 6.0.0) and the nf-core/nanoseq pipeline (version 3.1.0) (Chen et al., 2025) from the nf-core community (Ewels et al., 2020) was used to do all primary bioinformatic analyses. Briefly, this included the quality control (QC) of the reads with the nanoplot (version 1.41.6) (De Coster and Rademakers, 2023) and multiqc (version 1.9) (Ewels et al., 2016) programs and the mapping of the `fastq` files onto human and canine genomes using minimap2 software (version 2.15-r905) (Li, 2018). For the ten samples, we considered all reads having a Qscore >5 (**Table 1**). Intersection between all genomic features (i.e. TSS and CAGE data) was done using bedtools (version 2.27.1).

**Table 1.** Number of reads sequenced and aligned on canFam4 (CFA) and GRCh38 (HSA).

| Sample | Cellosaurus ID | Total Reads | Median Length | Primary Mapped | % Primary Mapped |
| --- | --- | --- | --- | --- | --- |
| CFA-MM1 | CVCL_OD14 | 4,002,891 | 806 | 3,408,273 | 85.15 |
| CFA-MM2 | CVCL_IZ11 | 8,843,341 | 769 | 6,253,908 | 70.72 |
| CFA-MM3 | Dog-OralMel-18395 | 7,108,351 | 943 | 5,901,302 | 83.02 |
| CFA-MM4 | Dog-OralMel-18333 | 6,877,092 | 896 | 5,696,283 | 82.83 |
| CFA-HS1 | Dog-HS-5472-lung | 11,177,946 | 826 | 9,157,969 | 81.93 |
| CFA-HS2 | Dog-HS-5472-spleen | 4,055,709 | 878 | 3,355,138 | 82.73 |
| CFA-HS3 | Dog-HS-7051 | 3,908,392 | 793 | 3,040,788 | 77.80 |
| CFA-OS1 | Dog-OSTEO-10188 | 6,816,094 | 1,050 | 5,707,582 | 83.74 |
| HSA-MM1 | CVCL_1282 | 5,457,058 | 886 | 4,865,343 | 89.16 |
| HSA-MM2 | CVCL_6797 | 4,734,796 | 852 | 4,025,056 | 85.01 |

### Benchmark analyses

#### Reference annotations and genomes

In the LRGASP study, only one reference annotation (Gencode) was used to benchmark tools in human LR-RNASeq samples. In our study, we bench-marked ANNEXA using several human and dog annotations in order to test the influence of the completeness of the input reference annotation in two species with high and medium quality reference annotations. We compiled reference genomes and annotations downloaded from Ensembl (Dyer et al., 2025), Refseq (O’Leary et al., 2016), Gencode (Mudge et al., 2025) or original papers (Varabyou et al., 2023; Wang et al., 2021) (**Table 2**).

**Table 2.** Species, assembly, reference annotation with gene and transcript counts. Links to the annotations can be found in **Supplementary Table 1**

| Species | Assembly | Reference Annotation (version) | Genes | Protein coding | lncRNA | Transcripts | Protein coding | lncRNA |
| --- | --- | --- | --- | --- | --- | --- | --- | --- |
| Dog | canFam4 | Ensembl (113) | 30,653 | 20,533 | 6,446 | 56,403 | 44,438 | 8,291 |
| Dog | canFam4 | UU (University Uppsala) | 29,535 | 20,072 | 9,177 | 158,561 | 143,888 | 13,476 |
| Dog | canFam4 | Refseq | 43,427 | 21,241 | 13,404 | 103,619 | 68,562 | 25,558 |
| Human | GRCh38 | Genecode (47) | 86,402 | 23,258 | 36,524 | 412,034 | 100,025 | 195,184 |
| Human | GRCh38 | CHES (3.0) | 63,755 | 22,083 | 18,598 | 168,451 | 105,328 | 36,354 |

The Gencode v47 annotation leverages the Capture Long-Read Sequencing (CLS) pipeline, combining Nanopore and PacBio sequencing to identify over 132,000 novel human lncRNAs, resulting in higher transcript counts compared to other annotations (Kaur et al., 2024). The CHESS annotation, by contrast, builds on Human RefSeq and incorporates RNAseq data from the GTEx project, offering broad tissue representation but without RNA enrichment followed by long-read sequencing. For the canine genome, the annotation provided by the University of Uppsala (UU) integrates Nanopore and PacBio data from 40 canine tissues, providing a comprehensive long-read-based resource (Wang et al., 2021). Unlike UU, Ensembl and RefSeq annotations lack a standardised LR-RNASeq transcriptome dataset across canine assemblies, leading to variability in gene and transcript counts, as recently reviewed (Hoffmann et al., 2026a).

#### Transcriptome reconstruction benchmarking

ANNEXA was run using Bambu and StringTie on the eight canine samples (bam files) with the canFam4 genome assembly and the three reference annotations (Ensembl, RefSeq and UU), and on the two human samples with the GRCh38 assembly and CHESS or Gencode annotations. For the Bambu runs, single-exon transcripts were discarded because their inclusion interfered with the calculation of the Novelty Detection Rate (NDR) recommended by the tool, leading to unstable filtering behavior. In contrast, StringTie retains single-exon transcripts by default when they meet the minimum coverage threshold. The full and filtered extended annotations produced after the runs were filtered to only keep genes and transcripts expressed (raw count >0) in at least one sample and were compared to the initial reference annotations using the GffCompare tool version v0.12.6 (Pertea and Pertea, 2020) which provides sensitivity and precision metrics at the exon, transcript and locus/gene levels.

True Positives (TP), False Positives (FP) and False Negatives (FN) were defined as followed:

- TP: Reconstructed features (exons, transcripts, or loci) that match corresponding features in the reference annotation. At the transcript level, GffCompare considers TPs as full exon chain matches between predicted and reference transcripts, where all internal exons match and outer boundaries can differ by at most 100 bases.
- FP: Reconstructed features that are absent from the reference annotation. These may represent novel transcripts or potential artifacts.
- FN: Features present in the reference annotation but missing in the extended annotations, indicating transcripts that were not recovered.

From these classifications, we thus computed Precision as TP/(TP + FP) and Recall or Sensitivity as TP/(TP+FN).

#### Orthology analysis

Canine and human novel genes and transcripts were mapped on target genomes (GRCh38 and CanFam4, respectively) using the liftoff program (Shumate and Salzberg, 2021) with default parameters. Then, query elements were classified into three classes with respect to the target reference annotation (Gencode and Ensembl, respectively): unmapped (query gene not mapped to target genome), mapped unknownGenes (query gene mapping to intergenic regions) and mapped knownGenes (query gene mapping by 50% coverage and 50% sequence identity to known genes from target annotation *e.g.* Gencode for query dog gene and Ensembl for human query gene).

## Results

We produced LR-RNASeq data from both human (n=2) and canine (n=8) cancer cell lines (**Fig. 1.B**) and applied ANNEXA using different genome assemblies and reference annotations for each species in order to study the robustness of the pipeline in its ability to extend and to quality control these annotations.

### Effect of annotation sources for the identification of known and novel genes/transcripts

Analysis of known and novel genomic features (genes and transcripts) reconstructed by Bambu and StringTie (with their default parameters) revealed striking differences with respect to the tool used and the completeness of the input reference annotation (**Fig. 3**).

#### Known genes and transcripts

For the canine genome (canFam4), Bambu reconstructed more known elements than StringTie, consistently across all three reference annotations tested (Ensembl, Refseq, University of Uppsala (UU)), with this difference being the most noticeable for the UU annotation (21,416 genes and 106,378 transcripts reconstructed with Bambu, 15,163 and 59,422 with StringTie) (**Fig. 3.A**). For the human genome (GRCh38), Bambu also identified more known elements than StringTie across the two annotations (CHESS and Gencode) with this difference being most pronounced for Gencode (27,341 genes and 87,249 transcripts reconstructed with Bambu compared to 18638 and 50,255 respectively with StringTie) (**Fig. 3.B**). Compared to dogs, the relatively low percentage of features reconstructed from a reference annotation in human (for example, only 21% of Gencode transcripts are reconstructed by Bambu) could be explained by the fact that human reference annotations are more exhaustive than in dogs and also because our experimental design contained more LR-RNAseq samples in dogs (n=8) versus human (n=2).

#### Novel genes and transcripts

For the canine genome (canFam4), StringTie consistently identified more novel elements than Bambu, independently of the three reference annotations tested (Ensembl, RefSeq and UU). However, we observed that this difference is most pronounced using the Ensembl annotation, where Bambu detected 3,134 novel genes and 3,600 novel transcripts compared to the StringTie set of genes and transcripts (12,671 and 20,184, respectively). Using the UU canine annotation yielded the lowest number of novel genes and transcripts identified by both StringTie (7,791 and 11,900) and Bambu (1,070 and 1,150), likely due to the greater initial isoform completeness of the UU reference annotation compared to Ensembl and RefSeq, thereby reducing the number of newly detected transcripts (**Fig. 3.C**). In the human genome (GRCh38), using the Gencode annotation as reference produced less novel elements than with CHESS with both Bambu and StringTie, as expected given the higher number of genes and transcripts catalogued in the latest version of Gencode (**Fig. 3.D**). Interestingly, we observe a difference in the novel gene reconstruction pattern between canine and human annotations, where StringTie reconstructs more genes in the three canine annotations and Bambu more in the two human annotations, despite StringTie consistently reconstructing more novel transcripts across all five annotations. This suggests that in a well annotated genome such as GRCh38, Bambu can still identify novel genes while StringTie primarily reconstructs novel isoforms of already known genes.

Regarding ANNEXA filtering operations, which evaluated the full-lengthness of reconstructed genomic elements (see **Methods**), we observed that it dramatically reduced the number of novel elements identified by both tools, but the decrease is more pronounced for StringTie. On average in dogs, 50% of Bambu and 16% of StringTie novel genes are being conserved after ANNEXA’s check for TSS validity, while in Humans 65% of Bambu genes and 46% of StringTie genes are conserved in CHESS, and 35% and 25% with Gencode. The proportion of filtered StringTie transcripts is more pronounced with Ensembl (dog) and CHESS (human) annotations, suggesting that these new transcripts may often be fragmented. Together, these findings highlight that reference annotation selection significantly impacts novel element discovery, with StringTie demonstrating particular sensitivity to this choice, especially in less well-annotated genomes such as canFam4. In addition, this shows that Bambu consistently demonstrated stronger performance in terms of precision and recall, particularly after applying ANNEXA’s filtering steps for gene and transcript full-lengthness (**Supp. Fig. 4**)

**Figure 4.**
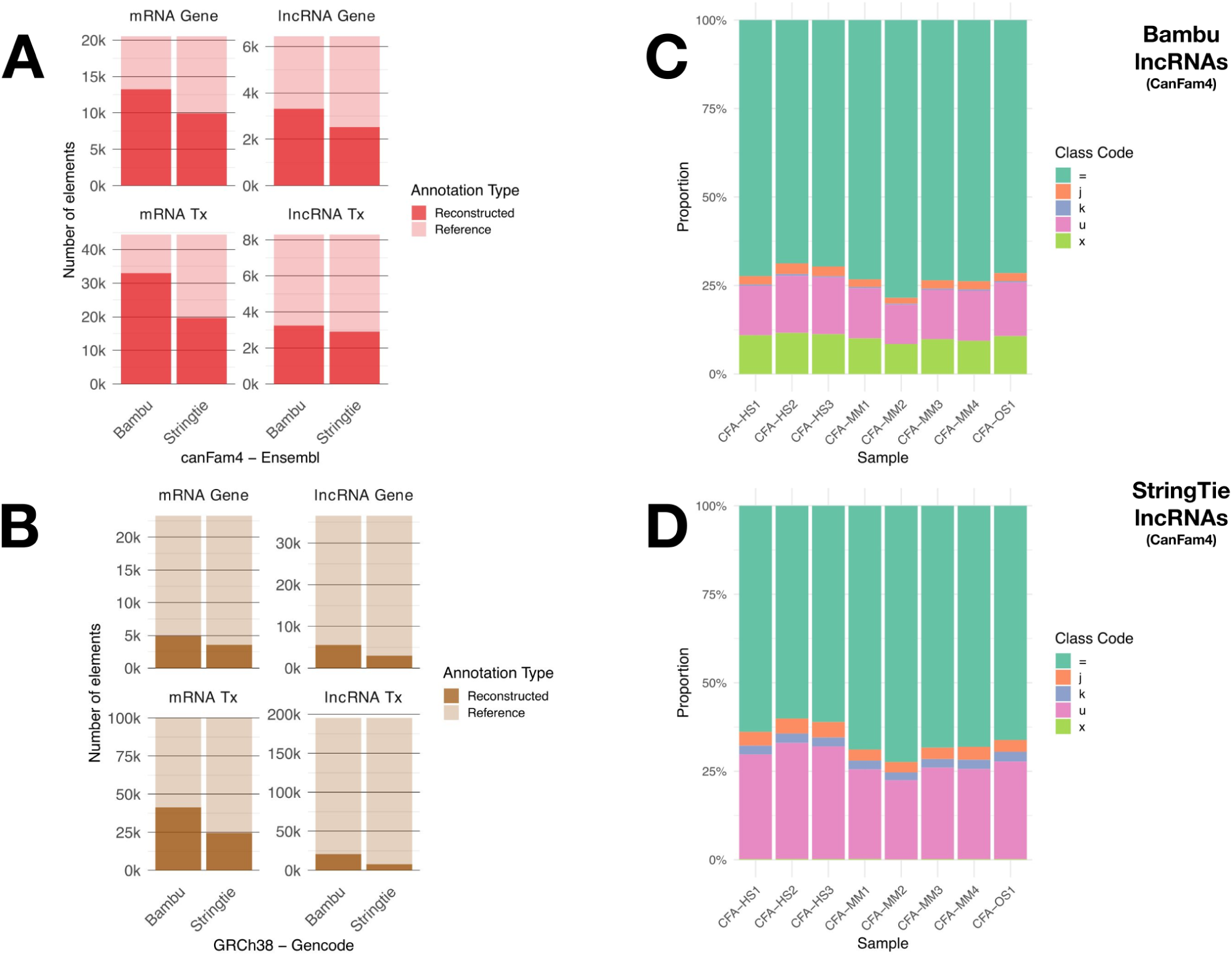
Annotation of lncRNAs/mRNAs by Bambu and StringTie. **A**- Number of known canine genes (top) and transcripts (bottom) for mRNA (left) and lncRNAs (right) annotated by Ensembl. **B**- Number of known human genes (top) and transcripts (bottom) for mRNA (left) and lncRNAs (right) annotated by Gencode. **C**- Class code distribution of canine lncRNAs annotated by Bambu. **D**- Class code distribution of canine lncRNAs annotated by StringTie. For panel C and D, class codes = (green): transcript isoforms matching the reference annotation, j (orange): novel spliced isoforms, k (purple): extension of reference gene, u (pink): intergenic transcripts and x (light green): antisense transcripts.

### Effect of species-specific reference transcriptome for the identification of known mRNAs and lncRNAs

Compared to protein-coding genes (mRNAs), long non-coding RNAs (lncRNAs) are considered particularly challenging to annotate due to their low expression levels and tissue specificity (Derrien et al., 2012; Kainth et al., 2023). As in our previous analyses, we evaluated the ability of Bambu and StringTie to reconstruct known genes and transcripts starting from long-read RNA sequencing (LR-RNASeq) data, this time distinguishing lncRNAs from mRNAs (**Fig. 4**).

In dogs (canFam4, Ensembl annotation), we observed that Bambu reconstructed a greater number of mRNAs and lncRNAs from the reference annotation than StringTie, with 13,239 vs. 9,930 genes for mRNAs and 3,314 vs. 2,525 for lncRNAs (**Fig. 4.A**). At the transcript level, we also noticed that the difference between Bambu and StringTie in term of reconstructed transcripts is more pronounced for mRNAs than for lncRNAs biotypes. A similar trend was observed in humans (GRCh38, Gencode annotation), where Bambu identified more genes and transcripts from the latest Gencode annotation than StringTie (**Fig. 4.B**). However, the proportion of reconstructed known genes was lower in humans than in dogs, again likely due to the smaller number of human LR-RNASeq data available as input.

Using ANNEXA’s transcript classification module, we then categorised both known (`class_code` `=`) and novel (all other `class_code`) mRNAs and lncRNAs annotated by the two tools from the eight canine and two human LR-RNASeq samples. Interestingly, this analysis revealed distinct patterns in the classification of novel canine lncRNAs: Bambu predominantly assigned them to `class_code` `u` (intergenic lncRNAs or lincRNAs, 14%) and `class_code` `x` (antisense lncRNAs, 10%), whereas StringTie primarily annotated novel lncRNAs as lincRNA (27%) and extension of known lncRNAs (2.4%), with very few classified as antisense (n=4) in dogs (**Fig. 4.C** and **Fig. 4.D**). This discrepancy is further illustrated in **Supp. Fig. 5**, where novel antisense lncRNAs are consistently detected by Bambu in both species but are largely absent in StringTie’s canine annotations, highlighting a tool-specific difference in antisense transcript reconstruction.

### Application of ANNEXA to mucosal melanoma data

Over the past decade, canine models have gained recognition as valuable spontaneous and immunocompetent systems for studying human cancers, particularly histiocytic sarcomas (HS) (Hédan et al., 2020) and mucosal melanoma (MM) (Prouteau and André, 2019). For MM, although rare in humans, it represents the most prevalent oral malignancy in dogs and exhibits notable clinical, biological, and genetic parallels with its human counterpart (Prouteau et al., 2022). Leveraging ANNEXA’s ability to balance precision and recall, we applied it to both canine and human data. In dogs, using a relaxed Bambu NDR threshold (NDR = 1) allowed to identify 9,612 novel genes (8,713 lncRNAs and 899 mRNAs) across the eight long-read RNAseq datasets (see **Methods**). Applying ANNEXA’s TSS validity filter significantly reduced the number of novel genes to 749 (595 lncRNAs and 154 mRNAs), indicating a likely high rate of false positives in the unfiltered set which could correspond to incomplete transcript models. Notably, the proportion of gene TSSs validated by orthogonal datasets, such as CAGE (Cap Analysis of Gene Expression) from the DogA consortium (Hörtenhuber et al., 2024), was significantly higher in the filtered set (52%, with 311 novel lncRNAs and 75 mRNAs validated) compared to the full dataset (9.3%, with 903 novel lncRNAs and 186 mRNAs validated). We also sought to assess whether novel genes in both humans and dogs could be conserved through evolution as a proxy for functional evidence (Ulitsky, 2016). We thus mapped the 9,614 novel canine genes (Ensembl - canFam4) on the human genome assembly used for the extended human annotation from Gencode and found 3,709 (38.6%) that could be mapped to GRCh38. Among these, 3,263 (88%) correspond to human genes from the Gencode reference annotation and thus represent novel orthologous relationships between novel dog and known human genes. Of these mapped genes, 2,901 (88.9%) were classified as lncRNAs, of which 931 matched with human lncRNA genes, while 362 (11.1%) were classified as protein coding and 240 of those were matching with human protein coding genes. Notably, we identified five novel canine lncRNA genes that also map to novel human genes (**Table 3**, **Supp.Fig. 6**>), with two of them being classified as protein coding in humans. Although these genes were not supported by CAGE data, they exhibited relatively high expression levels in both species, with mean read counts of 90.3 in dog and 66.1 in human MM samples. These conserved and expressed genes across species provide novel candidates for future functional validation and underscore the value of cross-species annotation to uncover novel, potentially functional elements.

**Table 3.**
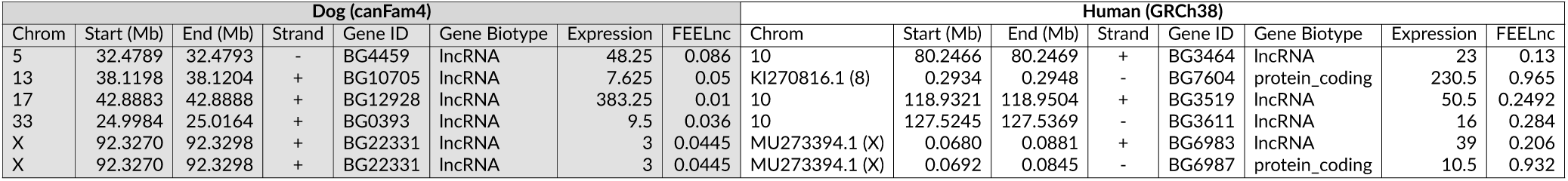
Genomic coordinates (in MegaBases, Mb), expression levels (raw counts) and FEELnc coding potential score of five novel orthologous genes identified in canine and human LR-RNASeq samples. Gene IDs have been shortened from “BambuGene” to “BG”.

| Dog (canFam4) |  |  |  |  |  |  |  | Human (GRCh38) |  |  |  |  |  |  |  |
| --- | --- | --- | --- | --- | --- | --- | --- | --- | --- | --- | --- | --- | --- | --- | --- |
| Chrom | Start (Mb) | End (Mb) | Strand | Gene ID | Gene Biotype | Expression | FEELnc | Chrom | Start (Mb) | End (Mb) | Strand | Gene ID | Gene Biotype | Expression | FEELnc |
| 5 | 32.4789 | 32.4793 | - | BG4459 | lncRNA | 48.25 | 0.086 | 10 | 80.2466 | 80.2469 | + | BG3464 | lncRNA | 23 | 0.13 |
| 13 | 38.1198 | 38.1204 | + | BG10705 | lncRNA | 7.625 | 0.05 | KI270816.1 (8) | 0.2934 | 0.2948 | - | BG7604 | protein_coding | 230.5 | 0.965 |
| 17 | 42.8883 | 42.8888 | + | BG12928 | lncRNA | 383.25 | 0.01 | 10 | 118.9321 | 118.9504 | + | BG3519 | lncRNA | 50.5 | 0.2492 |
| 33 | 24.9984 | 25.0164 | + | BG0393 | lncRNA | 9.5 | 0.036 | 10 | 127.5245 | 127.5369 | - | BG3611 | lncRNA | 16 | 0.284 |
| X | 92.3270 | 92.3298 | + | BG22331 | lncRNA | 3 | 0.0445 | MU273394.1 (X) | 0.0680 | 0.0881 | + | BG6983 | lncRNA | 39 | 0.206 |
| X | 92.3270 | 92.3298 | + | BG22331 | lncRNA | 3 | 0.0445 | MU273394.1 (X) | 0.0692 | 0.0845 | - | BG6987 | protein_coding | 10.5 | 0.932 |

## Discussion

In this work, we introduce ANNEXA, an all-in-one tool that not only performs transcriptome reconstruction but also simultaneously ensures quality control of extended annotations. Moreover, ANNEXA uniquely enables the comparative analysis of both novel mRNAs and lncRNAs derived from long-read RNAseq data, thereby enhancing their functional profiling and refining our understanding of their potential biological relevance. The comparison of two main transcriptome discovery tools and quantification methods reveals distinct trends depending on the reference annotation used. Our results show that Bambu consistently reconstructed more known genes and transcripts than StringTie, regardless of the reference annotation or the species (dog and human) analysed in this study. It also confirmed that the number of newly detected genomic elements is strongly influenced by the structure and completeness of the reference annotations.

The discrepancies between Bambu and StringTie could be attributed to their distinct methodological approaches. StringTie assembles transcripts by building splice graphs, without reference-based training, allowing it to retain a broader range of novel transcripts and include those that might deviate from known annotations. By contrast, Bambu uses a reference-guided machine learning approach to build a transcriptome, which prioritises precision by filtering out novel transcripts that are too dissimilar to the reference. This can explain why Bambu predicts fewer, higher-confidence transcripts, often limited to a single transcript per locus, while StringTie tends to predict multiple transcripts per locus, as observed in the Long-read RNA-Seq Genome Annotation Assessment Project (LRGASP) consortium’s study. It demonstrated that tools guided by a reference transcriptome, such as Bambu, perform optimally in well-annotated genomes and showed that Bambu predicted the fewest transcripts in de novo contexts, while StringTie predicted two or more transcripts for nearly half of the loci (Pardo-Palacios et al., 2024b).

The impact of reference annotation choice is also evident in the analysis of our human and canine long-read RNASeq data. As expected, using the latest Gencode release as reference produced fewer novel elements due to the higher number of genes and transcripts (86,402 and 412,034, respectively) already catalogued in its latest version. This underscores the importance of a comprehensive curated reference annotation to limit the detection of potentially artifact-driven elements.

To assess the impact of a more complete reference genome on transcript annotation, we also ran ANNEXA on the T2T-CHM13v2.0 assembly (Nurk et al., 2022) using two human mucosal melanoma long-read RNA-seq samples. ANNEXA identified 694 novel transcripts corresponding to 616 novel genes, including 54 genes overlapping CHM13-specific regions relative to GRCh38. In addition, novel isoforms from known genes were identified (**Supp. Fig. 7**), illustrating the benefit of combining complete genome assemblies with long-read transcriptome sequencing.

One challenge in benchmarking transcriptome reconstruction tools lies in the classification of predicted transcripts. Our evaluation focused on the recovery of features present in reference annotations, and following GffCompare’s documentation (Pertea and Pertea, 2020), we defined false negatives (FNs) as reference features absent from the extended annotations, and false positives (FPs) as novel features not present in the reference. In practice, some FPs may correspond to genuine, previously unannotated transcripts, while some FNs may represent transcripts not expressed in the analysed samples. Moreover, the exclusive use of reference annotations prevents the identification of true negatives, and thus precludes the computation of specificity. These limitations, which also apply for our TSS annotation with TransforKmers, highlight that performance estimates derived from reference-based benchmarking should be interpreted with caution, as they reflect both tool behaviour and the non-comprehensive nature of current annotations.

Using ANNEXA’s quality control module for known and novel long non-coding RNAs, we also observed this consistent trend where Bambu reconstructed a greater number of non-coding genes and transcripts than StringTie, both in dogs (canFam4, Ensembl) and humans (GRCh38, Gencode). However, the proportion of reconstructed known genes was lower in humans than in dogs for both tools, likely due to the more limited availability of human long-read RNA sequencing data in our study design, combined with the higher completeness of human reference annotations (Kaur et al., 2024).

Additionally, the classification of novel lncRNAs revealed distinct tool-specific patterns between Bambu and StringTie. Bambu primarily categorised novel lncRNAs as intergenic (lincRNAs) or antisense, reflecting its ability to detect antisense transcripts, even in contexts where reference annotations are incomplete, such as in canine samples. In contrast, StringTie assigned a higher proportion of novel lncRNAs to lincRNAs and novel isoforms of known lncRNAs, with only a few classified as antisense. This discrepancy is particularly pronounced in dog samples, where StringTie’s reference-guided assembly may limit its ability to reconstruct antisense transcripts. Conversely, Bambu’s approach appears less dependent on reference strand information, enabling consistent detection of antisense lncRNAs across species. These differences in classification outputs underscore the impact of transcriptome reconstruction tools and suggest that integrating multiple approaches may provide a more comprehensive representation of lncRNA diversity. Consequently, the choice of reconstruction tool should be guided by specific research objectives. For example, researchers focusing on natural antisense transcripts (NATs) in non-model organisms with incomplete reference annotation may benefit more from Bambu’s capabilities, whereas those aiming to extend known lncRNA annotations might prefer StringTie.

Our analyses further emphasises the need to carefully evaluate novel annotations generated by transcriptome discovery tools. The increase in detected genes and transcripts should be considered within the context of annotation robustness and biological validation. The LRGASP Consortium recommends incorporating additional orthogonal data and replicate samples when aiming to detect rare and novel transcripts. In our comparative oncology project, we used CAGE data from both dog and human samples to validate the full-length nature of novel transcripts. We found that only less than 10% of unfiltered novel transcripts (both coding and noncoding) were validated by at least one CAGE signal. Several factors could explain this relatively low proportion of TSS validation, including the fact that the CAGE sample conditions did not match those of our LR-RNASeq cancer cell lines and/or the earlier version of the Nanopore kit for library preparation (DCS109) could have led to truncated reads and, consequently, incomplete transcripts (Sessegolo et al., 2019). However, the use of ANNEXA’s module for TSS validation increased the proportion of validated genes to over 50%, highlighting the importance of combining complementary experimental and computational approaches to refine annotations.

In summary, this study highlights ANNEXA’s modularity in adjusting precision and sensitivity for accurate, context-specific coding and non-coding annotation. This flexibility is essential for subsequent experimental validation and for advancing our understanding of the role of the non-coding genome in biological processes.

## Availability and implementation

ANNEXA is written in Nextflow DSL2 (Di Tommaso et al., 2017). ANNEXA can run each process in conda environments as well as Docker or Apptainer containers, ensuring reproducibility and ease of use on different machines. The pipeline is available at https://github.com/IGDRion/ANNEXA. Input files and code to reproduce figures of this paper are also available here: https://github.com/IGDRion/ANNEXA/tree/main/Paper_Figures. The extended human and canine annotations from the results are available on Zenodo (DOI: 10.5281/zenodo.18195454).

## Acknowledgements

The authors thank the GenOuest bioinformatics core facility (https://www.genouest.org) for providing them with the necessary storage and CPU/GPU resources to perform the analysis, the IGDRIon facility for long-read sequencing, members of the emerging BINGO team (https://igdr.univ-rennes.fr/bingo-groupe-en-emergence) and BIS group (https://igdr.univ-rennes.fr/biologie-silico-0) for useful discussions. Preprint version xxx[change to the correct number] of this article has been peer-reviewed and recommended by Peer Community In XYZ[change to the name of the PCI] (https://doi.org/10.24072/pci.xxx [replace by the doi of the recommendation]; (**fake**)[replace by the citation of the recommendation]).

## Data, script, code, and supplementary information availability

All fastq files of the 10 LR-RNASeq samples are available through the ENA database under study ID: PRJEB88593. Script and codes are available online (https://doi.org/10.5281/zenodo.15209930; Hoffmann et al., 2026b). Supplementary information is available online (https://www.biorxiv.org/content/10.1101/2025.04.16.648718v1; Hoffmann et al., 2025).

---

**Supplementary Figure 1.**
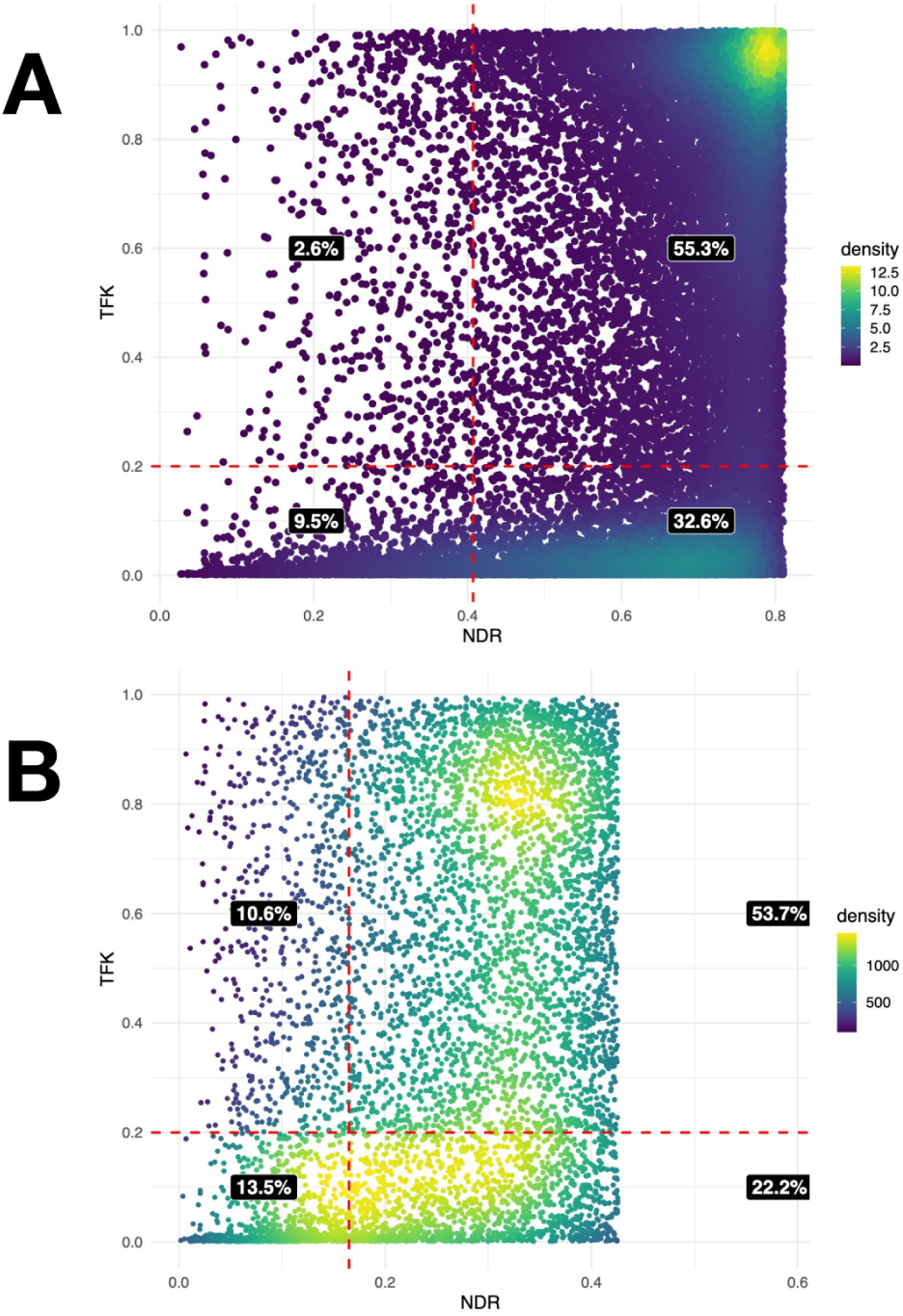
Distribution of novel transcripts identified by Bambu according to their TransforKmer score (TFK) and Novel Discovery Rate score (NDR) in canine samples with the Ensembl annotation (**A**) and human samples with the Gencode annotation (**B**). The red dotted lines represent the cutoffs used by ANNEXA to filter novel transcripts.

**Supplementary Figure 2.**
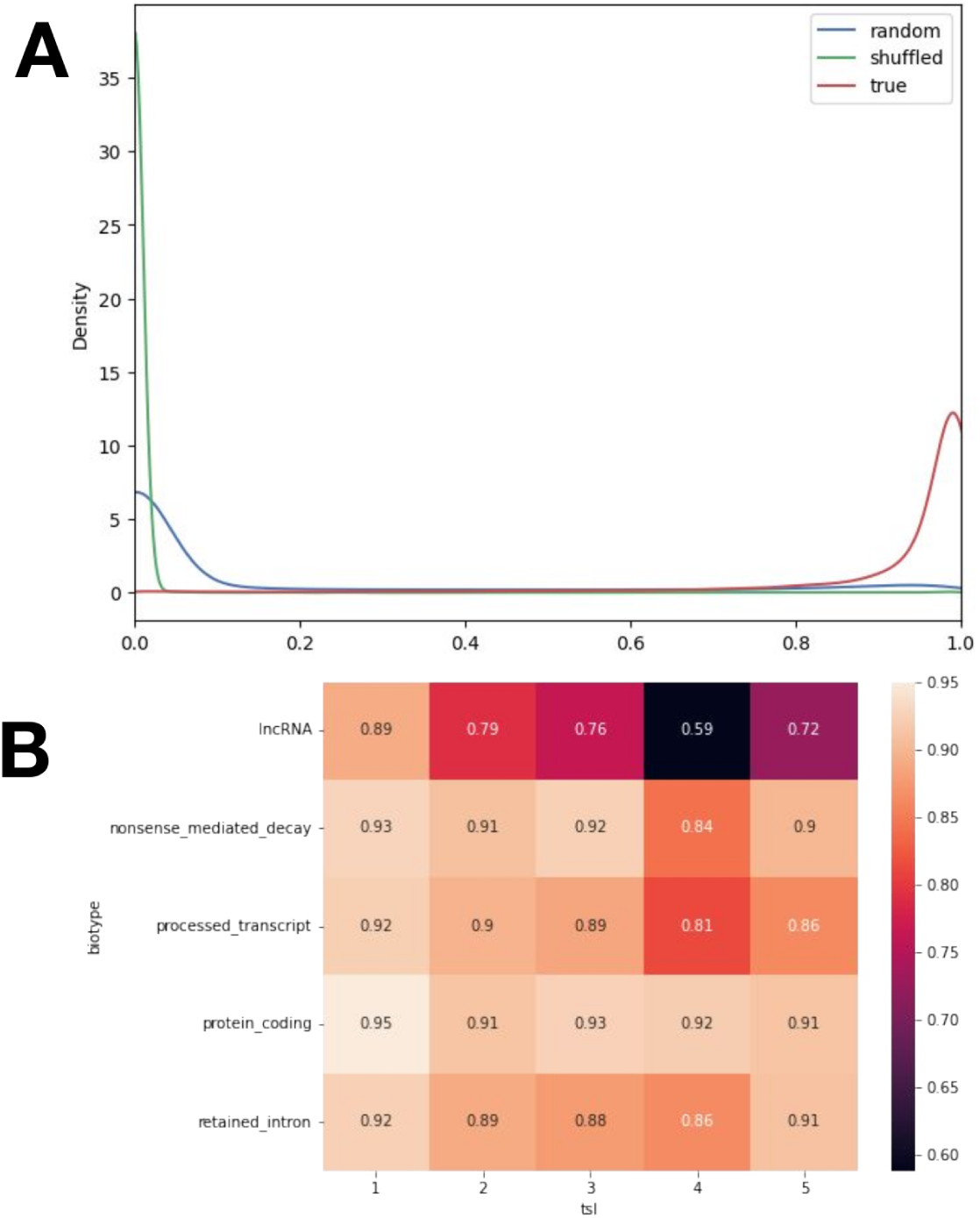
**(A)** Density plot of TransforKmers (TFK) prediction probabilities stratified by TSS sequence origins: random sequences (blue), shuffled (green) and true/reference TSSs (red). **(B)** Heatmap of TFK classification accuracy in human relative to transcript biotypes and transcript support level (tsl).

**Supplementary Figure 3.**
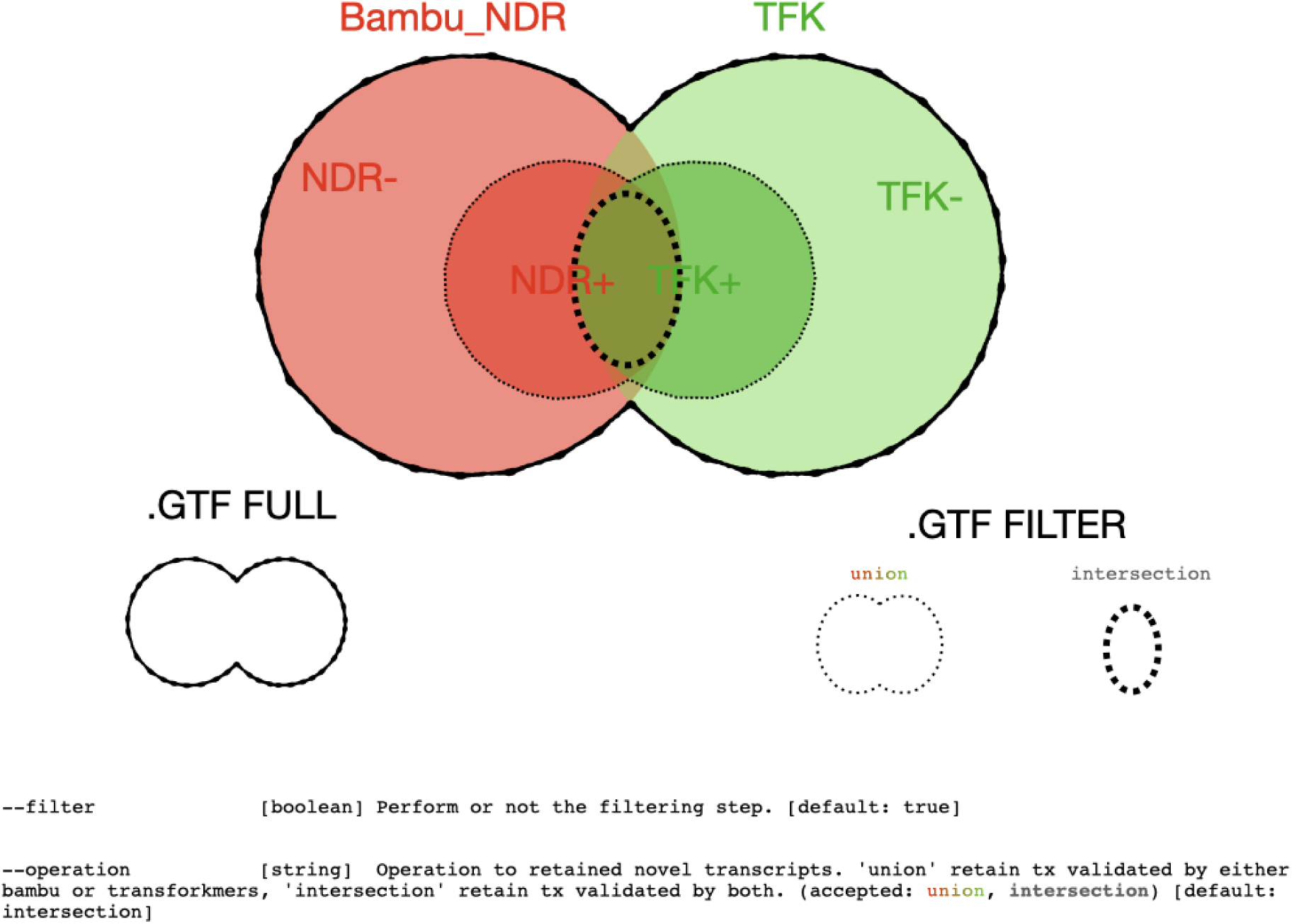
Filtering operations in ANNEXA. Novel transcripts can be filtered based on their NDR cutoff (in red, exclusive to Bambu) and/or TransforKmers (TFK) cutoff (in green, available with both Bambu and StringTie).

**Supplementary Figure 4.**
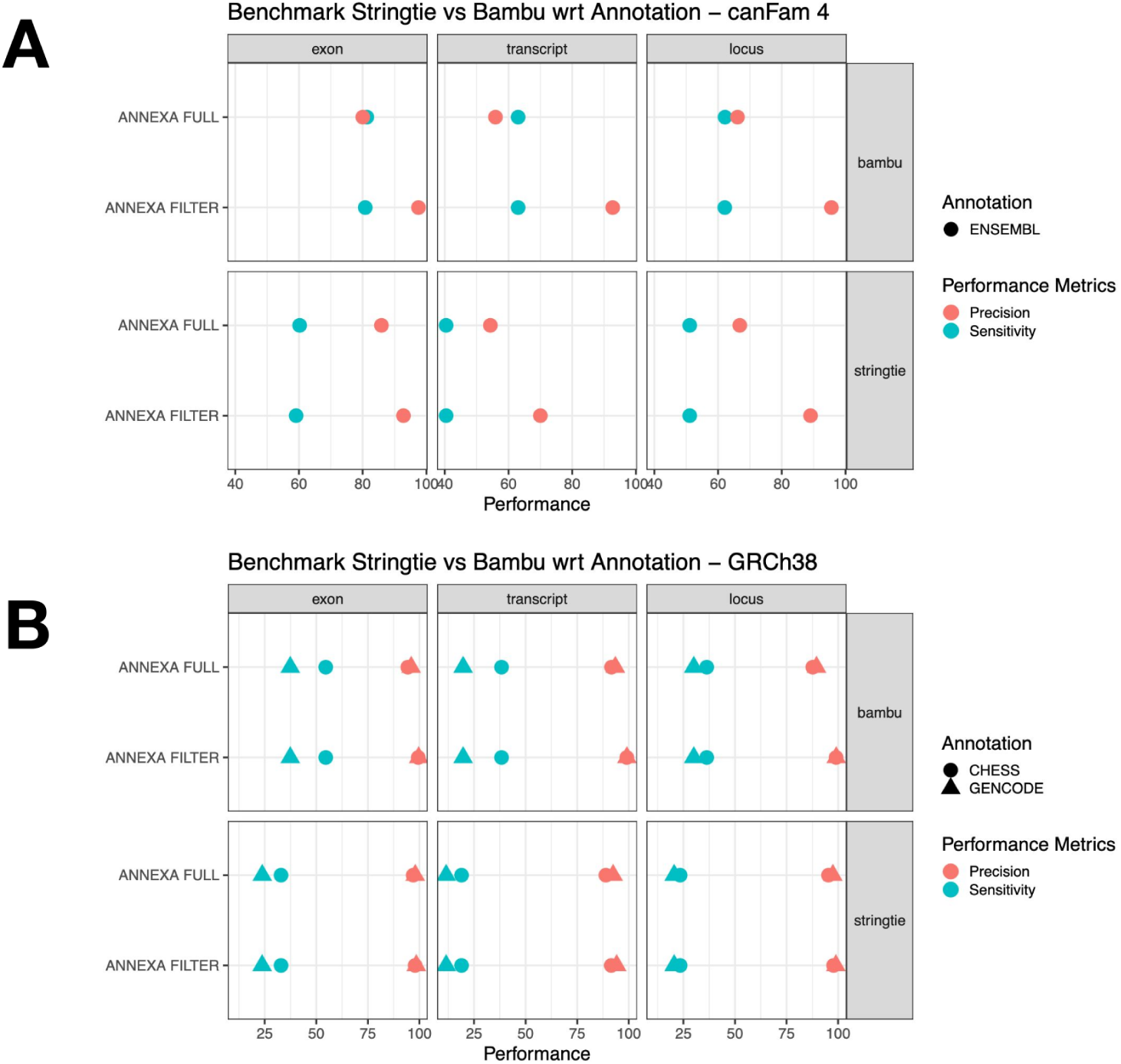
Comparison of precision (red) and sensitivity (green) metrics when using ANNEXA with Bambu or Stringtie on the canine (A) or human (B) LR-RNAseq data.

**Supplementary Figure 5.**
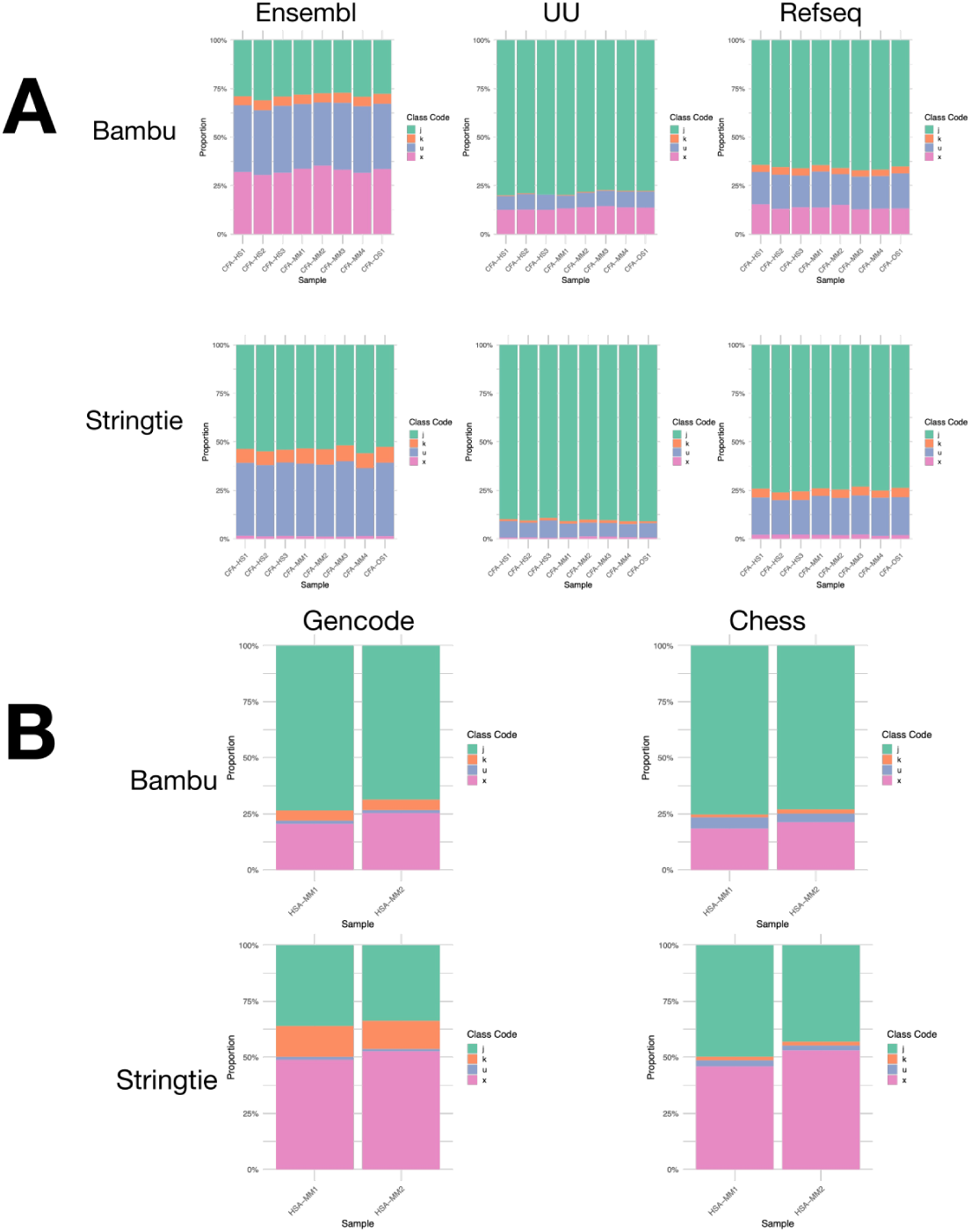
Classification of novel transcripts in dogs (**A**) and humans (**B**) with j = novel alternatively spliced transcript (green), k= extension of known transcript (orange), u = intergenic transcript (blue) and x = antisense transcript (pink).

**Supplementary Figure 6.**
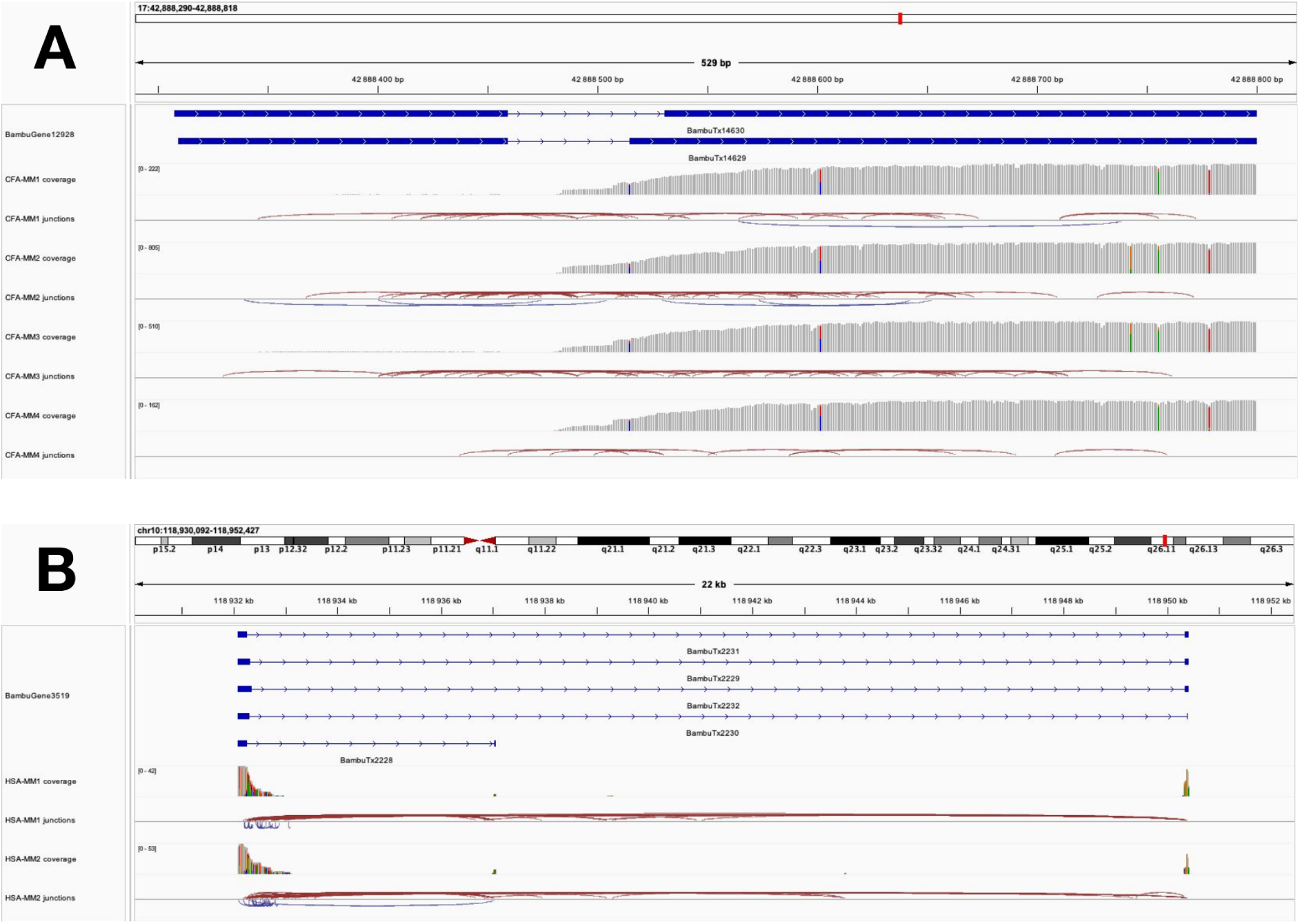
IGV screenshot of a novel orthologous gene both annotated in dog (A) and human (B). Long reads aligned from four mucosal melanoma (MM) samples in dogs (canFam4) and two MM samples in human (GRCh38) are represent below annotations.

**Supplementary Figure 7.**
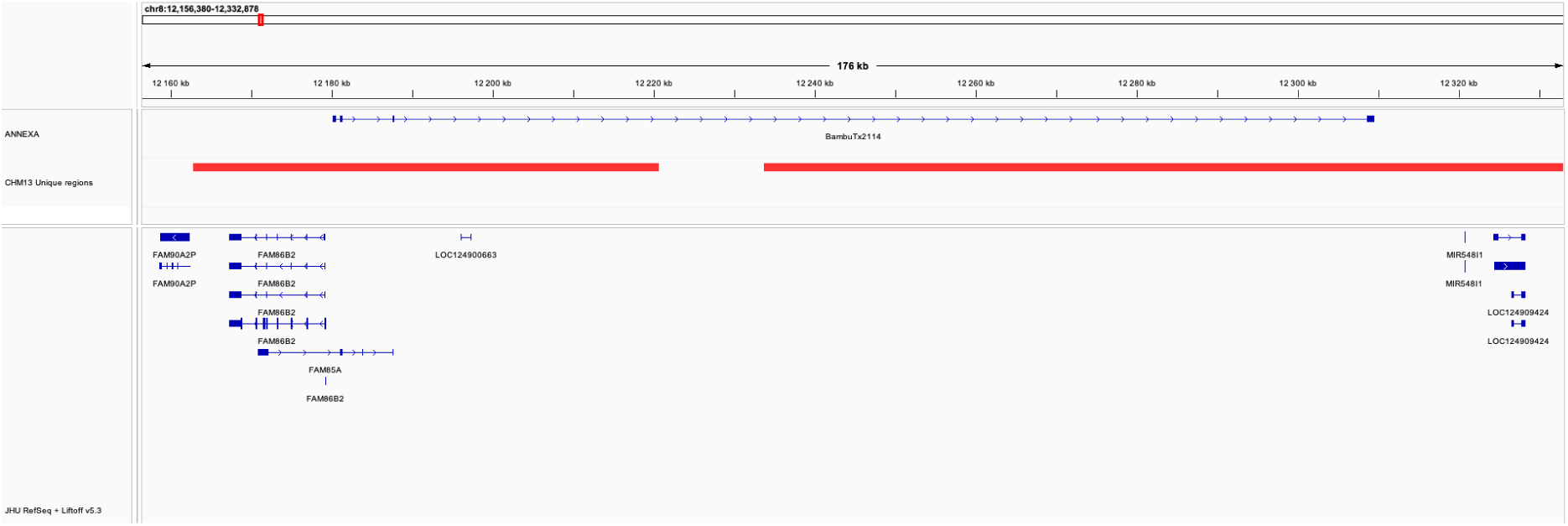
IGV screenshot of a novel isoform of the *FAM85A* gene (BambuTx2114), overlapping regions unique to CHM13 compared to GRCh38 (in red). In total, 54 novel genes and 93 novel isoforms of known genes were identified using the two human LR-RNAseq samples.

**Supplementary Table 1.** Links to canine and human annotations listed in Table 2.

| Species | Assembly | Reference Annotation (version) | Link |
| --- | --- | --- | --- |
| Dog | canFam4 | Ensembl (113) | https://ftp.ensembl.org/pub/release-113/gtf/canis_lupus_familiarisgds/ |
| Dog | canFam4 | UU (University Uppsala) | http://genome-euro.ucsc.edu/cgi-bin/hgTracks?db=canFam4 |
| Dog | canFam4 | Refseq | https://ftp.ncbi.nlm.nih.gov/genomes/all/GCF/011/100/685/ |
| Human | GRCh38 | Gencode (47) | https://www.gencodegenes.org/human/release_47.html |
| Human | GRCh38 | CHESS (3.0) | https://github.com/chess-genome/chess |

